# Multiplexed proteomics and imaging of resolving and lethal SARS-CoV-2 infection in the lung

**DOI:** 10.1101/2020.10.14.339952

**Authors:** Marian Kalocsay, Zoltan Maliga, Ajit J. Nirmal, Robyn J. Eisert, Gary A. Bradshaw, Isaac H Solomon, Yu-An Chen, Roxanne J. Pelletier, Connor A. Jacobson, Julian Mintseris, Robert F. Padera, Amanda J. Martinot, Dan H. Barouch, Sandro Santagata, Peter K. Sorger

## Abstract

Normal tissue physiology and repair depends on communication with the immune system. Understanding this communication at the molecular level in intact tissue requires new methods. The consequences of SARS-CoV-2 infection, which can result in acute respiratory distress, thrombosis and death, has been studied primarily in accessible liquid specimens such as blood, sputum and bronchoalveolar lavage, all of which are peripheral to the primary site of infection in the lung. Here, we describe the combined use of multiplexed deep proteomics with multiplexed imaging to profile infection and its sequelae directly in fixed lung tissue specimens obtained from necropsy of infected animals and autopsy of human decedents. We characterize multiple steps in disease response from cytokine accumulation and protein phosphorylation to activation of receptors, changes in signaling pathways, and crosslinking of fibrin to form clots. Our data reveal significant differences between naturally resolving SARS-CoV-2 infection in rhesus macaques and lethal COVID-19 in humans. The approach we describe is broadly applicable to other tissues and diseases.

**Summary:** Proteomics of infected tissue reveals differences in inflammatory and thrombotic responses between resolving and lethal COVID-19.

## INTRODUCTION

The great majority of tissue specimens acquired via biopsy, necropsy and autopsy in research and clinical settings are subjected to formalin-fixation and paraffin-embedding (FFPE). Molecular analysis of FFPE specimens is challenging but the use of fixed tissue has many advantages: FFPE samples are routine in diagnostic pathology and many archival samples are available, fixation better preserves tissue architecture than freezing, fixed tissues can be stored and shared easily, and use of thin sections (typically 5µm thick) conserves precious samples. In the context of COVID-19, fixation has the additional advantage that formaldehyde inactivates virus. Discovery proteomics of FFPE tissue is particularly difficult^1^ due to chemical modification of peptides and inhibition of sample proteolysis by crosslinking. A decade ago it was demonstrated that performing mass spectrometry (MS) on FFPE specimens is feasible^2^ and more advanced MS has recently been used to profile unfixed^3^ and FFPE tumor banks^4^ and also to assay sub-regions of a single specimen^5^.

SARS-CoV-2 infects cells that express angiotensin-converting enzyme 2 (ACE2) in concert with host proteases such as transmembrane serine protease 2 (TMPRSS2) and many of these cells are found in the respiratory tract (e.g. type II pneumocytes)^6^. Infection can cause uncontrolled activation of the immune system resulting in severe pulmonary injury and even death^7^, particularly in individuals with co-morbidities such as obesity, diabetes and hypertension. Study of the underlying molecular mechanisms has focused on bulk and single-cell RNA sequencing of relatively easy to access specimens such as blood, throat swabs and bronchoalveolar lavage^8^ and proteomics of serum^9^ and cell lines^10,11^. In humans, these data reveal strong but aberrant induction of interferon and inflammatory signaling^8,12,13^, upregulation of complement cascades^14,15^, and activation of platelets^16^.

In this paper we analyze detailed molecular and cell-level data on signaling pathways activated by SARS-CoV-2 infection using lung tissue obtained via necropsy of infected non-human primates (NHPs; rhesus macaques)^17^ in which infection naturally resolves and autopsy of humans who died from COVID-19. Acquiring these data required development of a proteomic method for protein extraction from multiple FFPE samples in a time series^18,19^ combined with multiplexed imaging^20^ of the same specimen. We accomplished this by developing a denaturing protein extraction protocol for FFPE samples involving thermal reversion of formaldehyde crosslinks. The extraction proteins were analyzed using stable-isotope tandem mass tag (TMT) MS in which individual sample digests are labelled with TMT, mixed together and then analyzed in a single MS run^21^. Key advantages of this approach are high accuracy of quantification (10% differences in protein abundance can be reliably detected)^18^ with few missing values across samples^22^. In the current work, capture of phospho-peptides on Fe-NTA^23^ following sample digestion and TMT labeling enabled quantification of protein phosphorylation. In human specimens, viral load could be measured by MS based on peptides derived from SARS-CoV-2 spike (S) and nucleocapsid (N) proteins; these peptides were not detected in rhesus macaque specimens, which contained substantially less virus. However, viral burden could be measured in animals using multiplexed cyclic immunofluorescence (CyCIF) imaging^20^.

The combined use of proteomics and imaging made it possible to characterize multiple steps in disease response from early cytokine accumulation and protein phosphorylation to activation of downstream receptor pathways, changes in cell state, and crosslinking of fibrin to form clots. Our data demonstrate interferon induction in NHPs that correlates positively with viral burden. In contrast, high viral burden with little interferon activation is possible in humans who died from COVID-19. Significant upregulation of cytokines, spatially widespread infiltration of macrophages and other immune cells, and thrombosis are observed in human decedents but are limited in animals to regions of lung injury. These and related findings exploit the strengths of multiplexed proteomics and imaging of intact tissue to provide a detailed molecular picture of naturally resolving SARS-CoV-2 infection and of lethal COVID-19.

## RESULTS

### FFPE proteomics of NHP and human specimens

We analyzed a cohort of SARS-CoV-2 infected adult NHPs, from which multiple specimens were collected 2, 4, or 14 days after infection as well as three animals re-challenged with virus 35 days after initial infection and necropsied 14 days later (Fig. 1a). NHPs are widely used to study viruses and vaccines^17,24–26^ and we and others have shown that SARS-CoV-2 infection results in robust viral replication in the upper and lower respiratory tracts followed by disease resolution within 7-14 days^17,27^. Multiple lung lobes were sampled from each animal and imaging showed that viral burden varied with days post infection and lung location^17^. This variation was particularly evident on day 2, at which point specimens with high and low viral loads were acquired from a single lung (Fig. 1b,c). For analysis by quantitative MS, nine specimens from infected and three from re-challenged rhesus macaques (n=8 animals) along with three specimens from an uninfected control animal were TMT-labelled and analyzed as a single mixed sample. Key findings were confirmed by repeat analysis of six additional day 2 or 4 post-infection specimens from the same animals (Extended Data Fig. 1a). Over ∼6,700 proteins were quantified per specimen and both principal component analysis (PCA) and two-way hierarchical clustering of MS data demonstrated separation by day of infection and viral load (Fig. 1d,e, Supplementary Tables 1,2). Interferon inducible genes (ISGs) co-clustered, as did proteins normally found in stoichiometric complexes such as subunits of the proteasome (Extended Data Fig. 1b,c). These data are consistent with retention of information on cellular pathways and macromolecular complexes during MS of FFPE samples.

**Figure 1.**
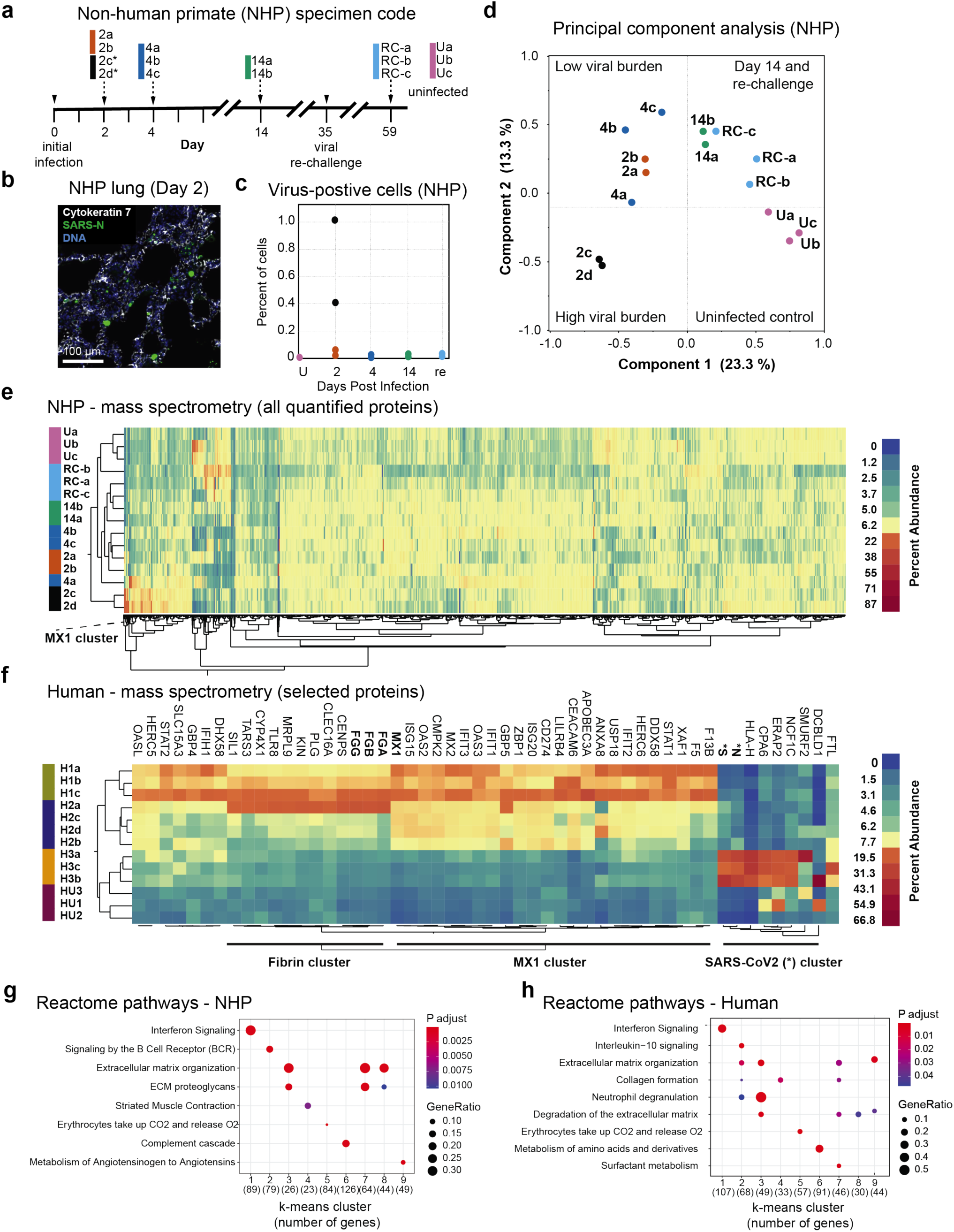
Formalin fixed paraffin embedded (FFPE) tissue proteomics of SARS-CoV-2 infected lung from non-human primate (NPHs) and human COVID-19 decedents. **a**, Specimen code for rhesus macaques (non-human primate; NHPs); details of the SARS-CoV-2 challenge schedule have been described previously.^17^ For specimens acquired two days post infection black (specimens 2a,b) and orange (2c,d) refer to regions of lung tissue with high and low viral burden as established by imaging - see panel b. **b**, Fluorescence micrograph of rhesus macaque lung 2-day specimen stained for cytokeratin 7 (grayscale), SARS nucleocapsid protein (SARS-N, green) and DNA (blue). Scale bar, 100µm. **c**, Quantification of virus positive cells in rhesus macaque lung sections by IF imaging; the denominator is all cells (nuclei) staining positive for SARS-N across the whole slide image. **d**, Principal component analysis (PCA) of quantitative mass spectrometry (MS) data. Resolution of uninfected and infected NHP samples over the trajectory of the disease mapped over components 1 and 2, as shown. **e**, Two-way hierarchical clustering of 6,767 quantified proteins in 15 NHP specimens over the course of infection; Specimens and color scheme as in a. Scaled relative protein abundance is shown from 0% (dark blue) to 87% (red). Position of the MX1 protein cluster is indicated. **f**, Subset of a two-way hierarchical clustering diagram for 6,729 quantified proteins from 10 human FFPE lung specimens (H1a-c, green; H2a-d, navy blue; H3a-c, yellow) and 3 uninfected (HU1-3, red) decedents; complete data are shown in Extended Data Fig. 1. The region shown here emphasizes clusters with MX1 (bold), fibrin (FGA, FGB, FGG; bold) and viral spike and nucleocapsid protein (S*, N*, bold). Scaled relative protein abundance shown from 0% (dark blue) to 66.8% (dark red). **g-h**, Reactome pathway enrichment analysis of the most variable proteins grouped based on k-Means clustering. The top pathway for each cluster is shown (FDR <0.05). In NHPs IFN signaling (cluster 1), B cell receptor signaling (cluster 2), complement cascade (cluster 6) and angiotensinogen processing (cluster 9) are also significantly enriched in separate cluster groups. In human decedents IFN signaling (cluster 1) and neutrophil degranulation (clusters 2 and 3) were significantly enriched. Protein ratios range from 0.1 to 0.3 in NHPs 0.1 to 0.5 in humans (scaled marker size). Pathway names have been abbreviated for clarity; complete data are shown in Extended Data Figure 2.

We also analyzed 13 human FFPE lung specimens collected at autopsy from three SARS-CoV-2 infected and three uninfected decedents (Supplementary Table 3)^28^. A similar number of proteins was quantified as in NHPs (Extended Data Fig. 1d and Supplementary Table 4) and co-clustering of ISGs (e.g. MX1), fibrins and clotting factors (e.g. Factors V, XIIIb and plasmin) was observed (Fig. 1f). The decedent with the highest viral levels of S and N viral proteins also had elevated levels of the angiotensin processing enzyme (CPA6), potentially reflecting binding of SARS-CoV-2 to angiotensin-converting enzyme-2 (ACE2)^29^. Reactome pathway analysis of rhesus macaque and human MS data (Fig. 1g,h, Extended Data Fig. 2a,b) revealed parallel enrichment of interferon, complement, immune-related, ECM and other pathways. As often observed in patients with severe COVID-19, decedents in our study had multiple pre-existing conditions (Supplementary Table 3). This complicates comparison of COVID-19 patients with each other and with controls. Nonetheless, in both monkey and human specimens, cluster analysis separated controls from SARS-CoV-2 infected specimens and pathway analysis revealed similar up and down-regulated pathways.

### Inflammation and its consequences

In NHPs, Gene Set Enrichment Analysis (GSEA; the term “gene sets” is used for simplicity but proteomic data were analyzed) revealed significant (FDR <0.05) upregulation of interferon type I and II (IFNα and IFNγ) pathways 2 and 4 days post infection compared to uninfected controls (Fig. 2a and Extended Data Fig. 3a,b). By day 14, when virus was undetectable^17^, IFN programs had largely returned to pre-infection levels. Activation of interferon pathways was confirmed by upregulation of multiple IFN-stimulated genes (ISGs) at days 2 and 4, with substantially stronger upregulation when viral load was high (compare NHP specimens 2a, b with 2c, d; Fig. 2b,c). In humans, the same IFN pathways were upregulated by viral infection, as were many of the same ISG proteins (Fig. 2b, right panel). In day 14 and re-challenged NHPs, the levels of ISGs had largely returned to normal but PCA and GSEA revealed differences relative to control animals; these differences largely involved pathways also upregulated at days 2 and 4 post infection.

**Figure 2.**
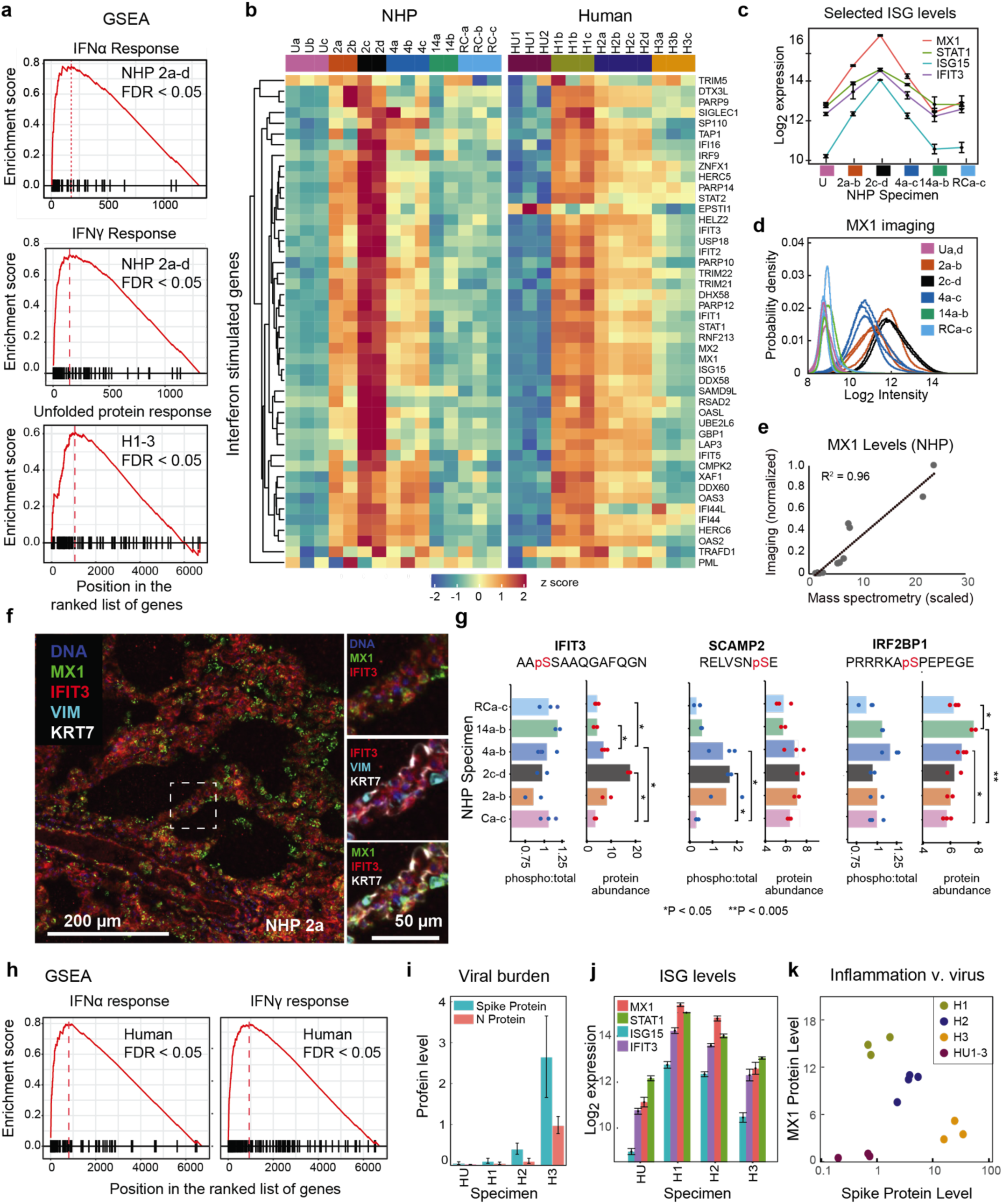
Analysis of Interferon-stimulated genes (ISGs) and anti-viral proteins from NHP specimens. **a**, Gene set enrichment analysis (GSEA; FDR <0.05) plot for signatures of IFNα and IFNγ in NHPs (all day 2 samples) and unfolded protein response in humans (all decedents) as compared to uninfected controls. The red plot corresponds to the running enrichment score for the pathway as the analysis walks down the ranked gene (protein) list, denoted by black bars on the X-axis; the location of the maximum enrichment score is shown. **b**, Comparison of levels of IFN-stimulated and anti-viral proteins (normalized TMT signal/noise) with z score ranging from −2 (blue) to 2 (red). Specimen color schemes as in Fig. 1a. **c**, Levels of selected INF-stimulated genes MX1 (red), STAT1 (green), ISG15 (blue), and IFIT3 (purple) in NHP samples. Values indicate mean +/− standard error **d**, Distribution of quantified single-cell MX1 staining intensity in cells from individual specimens using color code of Fig. 1a. **e**, Correlation of scaled MX1 protein levels measured using TMT-MS and quantitative IF microscopy. **f**, IF image of rhesus macaque lung sample 2a stained for MX1 (green), IFIT3 (red), vimentin (cyan), cytokeratin 7 (gray) and DNA (blue). Scale bar, 200µm. Inset region (dotted square) illustrates MX1 staining in nearly all vimentin-positive cells and IFIT3 in many such cells. Scale bar, 50 µm. **g**, Quantitative comparison of relative protein abundance (scaled) of IFIT3, SCAMP2 and IRF2BP and ratio of the indicated phospho-peptides (scaled) to the total protein abundance (scaled). Data for each sample and mean; *P <0.05, **P <0.005 with n=2-3. **h**, GSEA plot (FDR <0.05) for hallmark IFNα and IFNγ signatures in human decedents H1-H3 compared to uninfected samples. **i**, Levels of viral proteins quantified in human samples by MS; values denote mean +/-standard error of three specimens per decedent. **j**, Expression of selected IFN-stimulated genes MX1 (red), STAT1 (green), ISG15 (blue), and IFIT3 (purple) in H1-H3. Values indicate mean +/-standard error. **k**, Correlation of relative protein quantification for MX1 and SARS-CoV-2 spike protein in H1-H3.

Multiple significant changes observed in proteomic data were confirmed by immunofluorescence (IF). In NHPs, imaging showed that MX1 levels peaked at day 2 (Fig. 2d) and MX1 levels measured by IF and MS correlated well across all specimens analyzed (Fig. 2e). MX1 staining was observed in multiple types of immune cells, and punctate intracellular staining was consistent with assembly of MX1 into higher-order structures (Fig. 2f)^30^. MS also quantified changes in the phosphorylation of multiple proteins involved in IFN transduction and responses including IFIT3^31^, SCAMP2^32^, and IRF2BP1 (see Supplementary Table 5 for all quantified phosphosites). For IFIT3 and IRF2BP1, both protein and phospho-site levels increase in parallel and in other cases, SCAMP2 for example, the stoichiometry of phosphorylation increased in specimens with high viral burden; both patterns of modification reflect known regulation of IFN mediators (Fig. 2g). By imaging, IFIT3 levels were elevated in day 2 and 4 specimens as in proteomic data and exhibited a diffuse cytoplasmic pattern in epithelial, endothelial and some immune cells (Fig. 2f). In specimens from re-challenged animals, which are resistant to reinfection^17^, no significant activation of IFN pathways or induction of ISGs was observed. However, GSEA revealed enrichment for plasma and NK cells (Extended Data Fig. 3c), which is consistent with an antigen-specific and other anti-viral responses^33,34^. Overall these data demonstrate a classical IFN-mediated response to viral infection in NHPs that is proportional to viral burden. In human specimens, GSEA also revealed significant enrichment for IFNα and γ responses (Fig. 2h), but induction of ISGs was not obviously related to viral load (Fig. 2i,k): for example, decedent 3 with the highest viral load exhibited minimal ISG induction at death. Although we cannot exclude the confounding effects of co-morbidities in this small sample of decedents, our proteomic and imaging data are consistent with previous reports of an impaired or dysregulated IFN response in COVID-19 patients^8,35^.

Additional, significantly upregulated pathways in human decedents and high-virus NHP specimens included the unfolded protein response and mTORC1 signaling (Fig. 2a, Extended Data Fig. 3d). The former is consistent with the hypothesis that SARS virus infection induces proteostatic stress^36^ and might be considered for therapeutic intervention^37^. mTORC1 activation has been linked to insulin resistance and induction of type II diabetes in hepatitis C virus infection^38^. A similar association has also been suggested for COVID-19: new-onset diabetes is observed with COVID-19^39^ and diabetes is a known co-morbidity^40^. Our data are consistent with a mechanistic link between SARS-CoV-2 infection and the etiology of type II diabetes^41^.

### Cytokines and immune activation

Severe COVID-19 is associated with dramatic increases in cytokine levels^13^ and cytokine storms are postulated to play an important role in disease pathology^42^. Measuring cytokines *in situ* in tissue has historically been difficult, but we quantified 35 cytokines in human and 13 in macaque samples by MS. In one or more human decedents we found more cytokines significantly upregulated (11; P < 0.05) or downregulated (5) than in NHPs (Fig. 3a,b and Extended Data Fig. 4a). Specimens from human patient 2 (H2) exhibited a particularly strong induction of multiple cytokines: IL-1β, CXCL1/5/10 and CCL19/20 were all significantly upregulated (P < 0.05) as were protein components of the NLRP3 pathway (Fig. 3a,b). The NLRP3 inflammasome is a key component of the innate immune system that regulates secretion of pro-inflammatory cytokines IL-1β and IL-18 in response to infection and cellular damage^43^. Upregulation of IL1RAP (Fig. 3a, bar plot), a co-receptor for the IL1 receptor type 1, suggests enhanced responsiveness to these cytokines. ssGSEA also revealed enrichment of pathways regulated by IL-10 and IL-1 along with TNFα signaling via the NF-κB pathway (Fig. 3c). IL-6 and IL-10, which are known to play a role in severe COVID-19^44,45^, were not quantified in our proteomic data, but upregulation of IL-6 and IL-10 response genes was evident by ssGSEA in two of three patients (Fig. 3d). We surmise that both interleukins escaped detection in our MS runs but might be quantifiable using targeted MS. While dysregulation of NLRP3^46^ and NF-κB^35^ has previously been described in severe COVID-19 cases based on whole-blood transcriptomics and cytokine profiling, our data demonstrate that this occurs in infected lung tissue and is associated with changes in the levels of cytokines, receptors and downstream response proteins at the site of infection.

**Figure 3.**
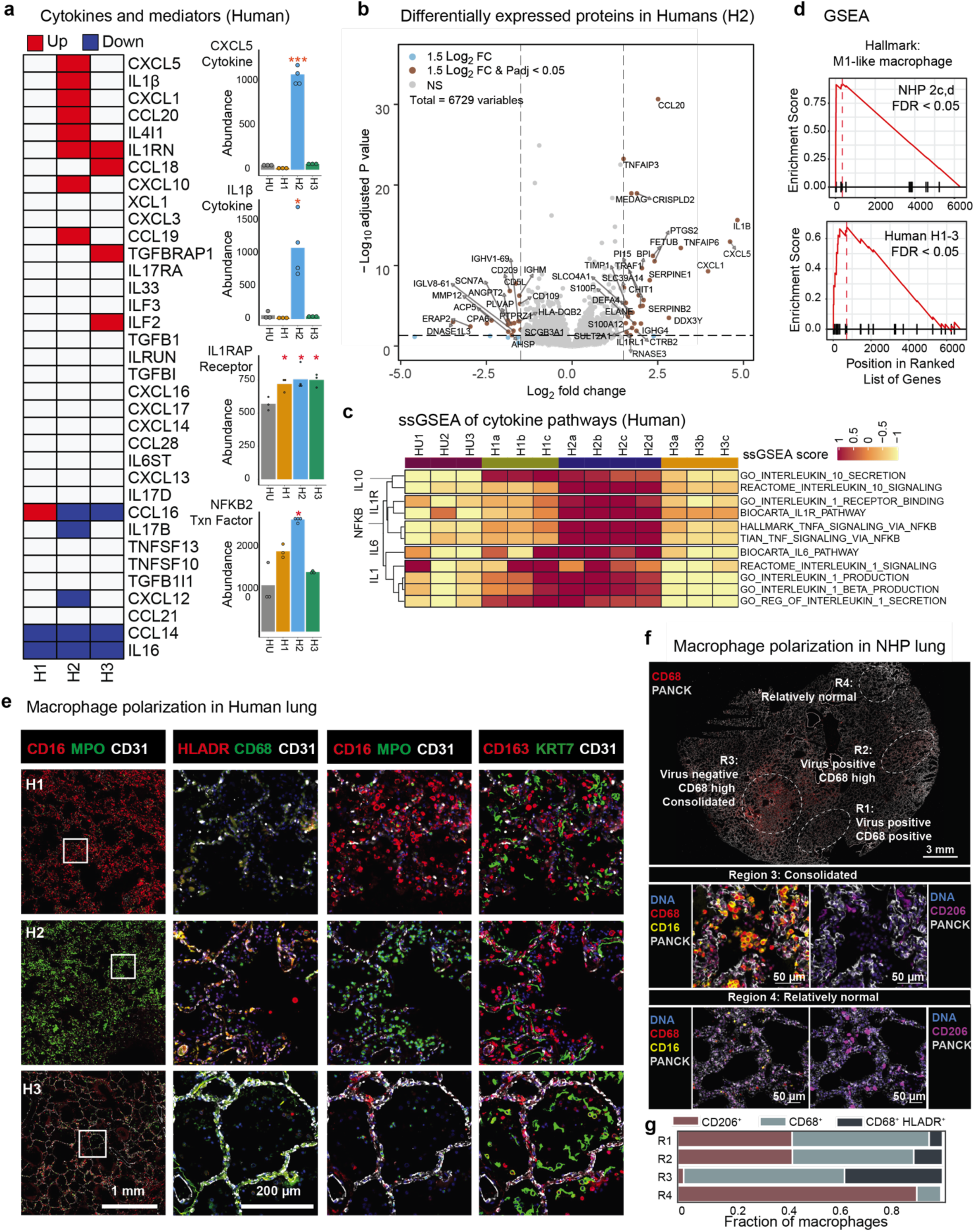
Cytokine levels and inflammation in lung tissue from human decedents. **a**, Binary heat map (left) showing significant increase (red; P < 0.05, t-test) or decrease (blue) of cytokines as quantified by MS in H1-H3. Bar plots (right) for selected cytokines showing mean protein abundance in control and human decedents (*P < 0.05, ***P < 0.001). **b**, Volcano plot of fold-change (samples H2a-c compared to HUa-c) for relative quantification of 6,729 proteins by TMT-MS. Shown are proteins with >1.5-fold Log2 fold-change without (blue dots) or with FDR <0.05 (brown dots), and proteins exhibiting no statistically significant change in abundance (grey dots). **c**, Single-sample GSEA (ssGSEA) heatmap showing enrichment of IL-1, IL-1R, IL-6, IL-10 and NF-κB related gene signatures in H1-H3. Enrichment scores are normalized by the absolute difference between minimum and maximum values and scaled between −1 and 1. **d**, GSEA plot for M1 like macrophage signature (FDR <0.05). Upper panel: NHP 2c,d vs. NHP Ua-c; lower panel human H1-H3 vs.HU1-3. **e**, CyCIF images of human lung parenchyma from H1-H3 stained for CD16 (red) MPO (green), CD31 (gray). Scale bar, 1mm. Alveolar regions (inset) are stained for cytokeratin 7 (green), CD163 (red) and CD31 (gray) to illustrate epithelial shedding and macrophage recruitment. Patient samples differ in staining for CD16 (red), MPO (green) and exhibit little or no cellular staining for CD68 (green), HLA-DR (red). Scale bar, 100µm. **f**, CyCIF images of NHP day 2 lung parenchyma stained for CD68 (red) and pan-cytokeratin (PANCK; grey). Scale bar 3mm. Regions of viral infection, consolidation and relatively normal morphology are highlighted (as scored by CyCIF and H&E staining of adjacent sections) with magnified views shown below showing staining for immune cell markers CD68 (red) CD16 (yellow) and CD206 (magenta). Tissue architecture is shown by DNA (blue) and PANCK (grey). Scale bar, 50µm. **g**, Percent stacked bar plot depicting the proportion of CD68^+^ and CD68^+^ HLADR^+^ inflammatory macrophages versus CD206^+^ M2-like macrophages (CD206^+^ cells) in virus-infected regions R1 & R2, consolidated region R3 and relatively normal lung R4.

Immune cells shape and respond to the cytokine microenvironment. When ssGSEA was performed on human proteomics data, we observed enrichment for multiple myeloid cell lineages with patterns of infiltration varying by decedent (Extended Data Fig. 4b). By GSEA, significant enrichment of an M1 macrophage^47^ gene signature and other inflammatory pathways was observed in human specimens (Fig. 3d, lower panel). To confirm this, CyCIF staining was performed on adjacent tissue sections using antibodies against CD16 (an NK and myeloid marker), CD163 (a macrophage marker), cytokeratin 7 (which labels alveolar epithelia), CD31 (a label for blood vessels and the alveolar wall) and myeloperoxidase (MPO; a neutrophil marker also expressed in monocytes, granulocytes and macrophages). This revealed distinct immune cell types in each human specimen (Fig. 3e). H1 primarily contained CD16^+^ CD163^+^ macrophages (red in left panels) whereas H2 contained CD163^+^ MPO^+^ macrophages (green in left panels; these cells were also CD68 and HLA-DR negative), consistent with activation of distinct classes of inflammatory macrophages in the two patients, neither of which exhibited a classical M1-like marker distribution. Remarkably, lung specimens from H3, which had the highest viral burden, contained few if any immune cells and only few enriched immune pathways by ssGSEA; H3 also declined rapidly after hospital admission (Fig. 3e; Extended Data Fig. 4b and Supplementary Table 3). In NHPs, immune signatures were upregulated primarily at day 2 in virus-high samples (“2c-d”, Extended Data Fig. 4c) as was an M1-like macrophage signature (FDR <0.05, Fig. 3d, upper panel). Imaging of serial sections confirmed recruitment of CD16^+^ CD68^+^ macrophages to regions of tissue damage (consolidated lung regions, Fig. 3f,g)^17^ consistent with a classic inflammatory phenotype. Together these data demonstrate strong activation of innate immune responses in both human and NHP specimens, with considerable diversity among human decedents.

### Complement activation and thrombosis

In contrast to other respiratory viruses, SARS-CoV-2 infection is associated with activation of complement cascades^48^ and microvascular thrombosis^49^. By ssGSEA, activation of both the classical and alternate complement pathways was detected in H1 and H2 (but not in H3); the lectin pathway was downregulated in all patients as compared to controls (Fig. 4a,e). By MS, we quantified levels of fibrin Q256-K437 crosslinked peptides (Fig. 4b), a major form of fibrin α-chain crosslinking previously characterized by MS using purified components^50^. Q256-K437 crosslinking was proportional to α-chain abundance (Fig. 4c). These data demonstrate factor XIII transglutaminase activation, the endpoint of the blood clotting cascade. Imaging also confirmed the presence of thrombi within the lung microvasculature (Fig. 4d). In H2 we observed upregulation of proteins involved in fibrin activation (e.g. prothrombin, thrombin, tissue factor, factor XI) as well as inhibition of clotting suggesting homeostatic resolution of thrombi. Control patient HU2 died of an acute myocardial infarction and exhibited significant enrichment for pathways related to activation of platelets, but neither to fibrin nor complement; the thrombotic signatures in HU2 were therefore distinct from those of COVID-19 decedents (Fig. 4e).

**Figure 4.**
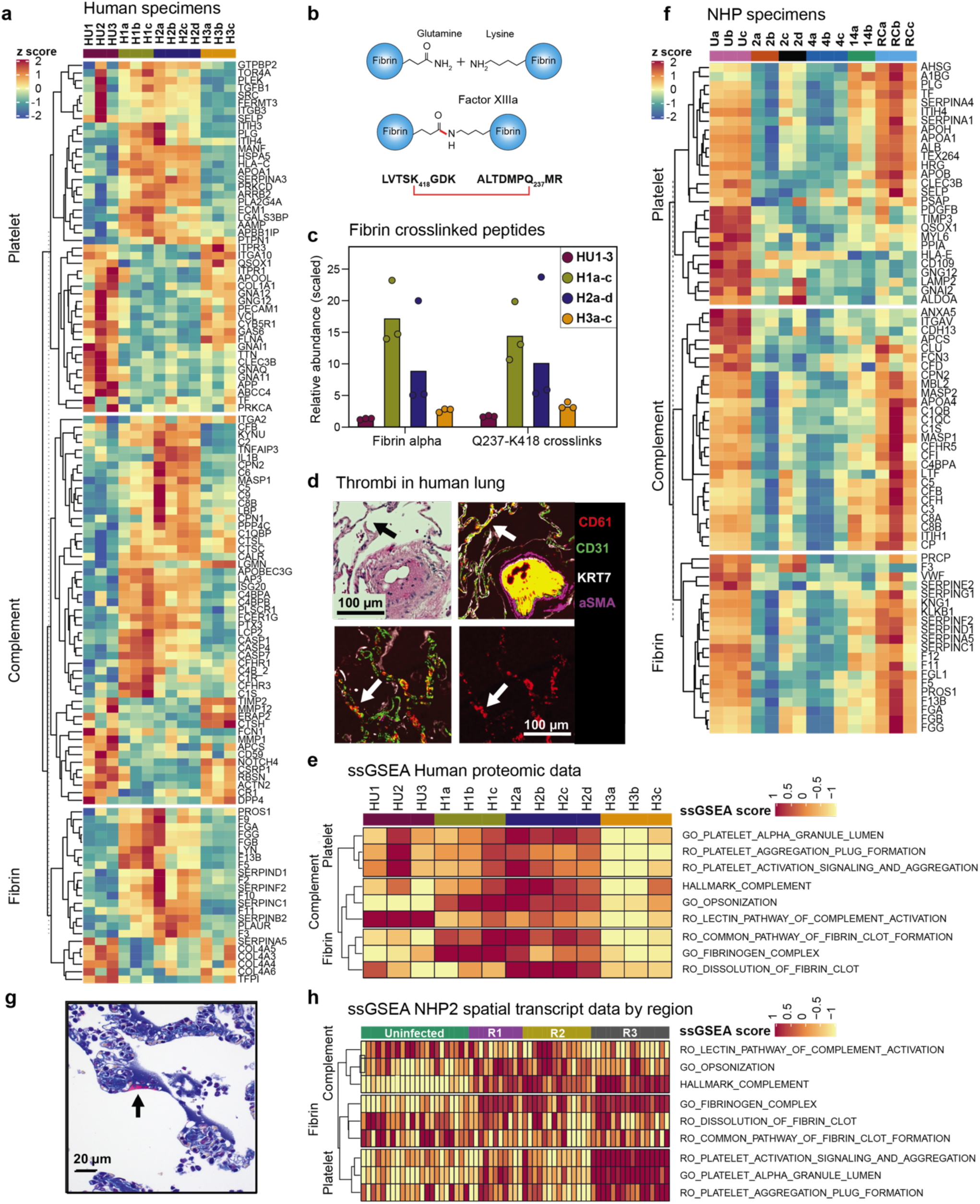
Analysis of complement activation and microvascular thrombosis in FFPE lung samples from human decedents and NHPs. **a**, Heatmap of human specimens showing the row scaled expression (z-score; from −2 (blue) to 2 (red)) of proteins involved in thrombosis and the complement pathway. Columns represent individual samples grouped by patients and each row corresponds to an individual protein. The dendrogram represents hierarchical clustering using Euclidean distance. **b**, Schematic of factor XIIIa transglutaminase-catalyzed fibrin cross-linking between lysine 418 and glutamine 237 of fibrin-α. **c**, Relative levels of total fibrin alpha (blue) and Q237-K418 cross-linked fibrin alpha (red) in human specimens. **d**, H&E (upper left) and CyCIF images (upper right) of blood vessel thrombosis in decedent C2 and capillary thromboses in HP3 (bottom right and left) stained for CD61 (red), CD31 (green), cytokeratin 7 (gray), and smooth muscle actin (magenta). Capillary thromboses are denoted with arrow. Scale bar, 100µm. **e**, ssGSEA heatmap showing the enrichment of platelet, fibrin and complement related gene signatures in human decedents. Enrichment scores are normalized by the absolute difference between minimum and maximum levels and scaled between −1 to 1; dendrogram represents hierarchical clustering using Euclidean distance. Reactome is abbreviated to RO in signature names. **f**, Analysis for NHP samples as in a. **g**, Carstairs’ stain of capillary thromboses in day 2 NHP specimen (2c); arrow denotes fibrin deposition along hyaline membrane. **h**, ssGSEA of spatial RNA transcription data (GeoMx) data from uninfected NHP animals and a virus-high day2 specimen. Morphological regions R1-R3 are shown in Fig. 2f.

Unexpectedly, in NHP samples, we observed significant downregulation (FDR <0.05) of complement, fibrin, and platelet-related signatures early in infection with recovery by day 14. Capillary thrombi were limited in extent and most apparent in day 2 specimens (Fig. 4f,g). To better understand this phenomenon, we performed spatial transcriptomic analysis with the GeoMx DSP platform^51^ on uninfected lung (22 regions) or infected lung with different levels of inflammation (16 high, 14 mid, and 11 low from the specimen in Fig. 3f). Regions of lung with high viral load but lacking overt tissue damage exhibited activation of complement pathways, whereas consolidated regions had upregulation of platelet aggregation, fibrin related pathways and downregulation of fibrinolysis (Fig. 4h). We therefore propose that in NHPs complement pathways and platelets are locally active during acute viral replication and consequent tissue consolidation and damage, as they are in humans. As disease resolves, homeostatic regulation decreases clotting and in NHPs this manifests as a general downregulation of signatures for complement, fibrin and platelets.

## DISCUSSION

In NHPs, SARS-CoV-2 infection is characterized by acute IFN induction proportional to viral burden as well as local induction of thrombotic programs. As virus is cleared over a two-week period, lung physiology returns to normal^17^. In contrast, severe COVID-19 in humans causes IFN pathway dysregulation, accumulation of immune cells^8,13,52,53^, microvascular injury, and thrombosis^54^ and we detect these pathologies by tissue proteomics and imaging. Significant increases in cytokine levels are evident in some human decedents but IFN induction is not in proportion to viral burden, with the decedent whose viral burden was highest exhibiting the weakest IFN response. The immune infiltrates, including macrophage activation states, also differ among decedents. Strong upregulation of pathways involving complement, fibrin processing and platelet activation is associated with extensive fibrin crosslinking and formation of thrombi. Because SARS-CoV-2 responses in NHPs are similar to those previously described for other respiratory viruses^55^, we propose that observed differences between infected NHPs and humans primarily arise from differences between naturally resolving infection and end-stage disease in the presence of co-morbidities.

Parallel analysis of fixed lung tissue by multiplexed mass spectrometry is a powerful counterpart to more common transcriptional and cytokine profiling of bronchoalveolar lavage and blood specimens in characterizing infection. The data in this paper highlight the ability of proteomics, imaging, and spatial transcriptomics of the same specimens to trace successive steps in diverse regulatory cascades. We quantify changes in cytokine levels, cognate receptors, signaling proteins, and downstream consequences of enzyme activity (e.g. phosphorylation, fibrin crosslinking), cell state and function. To our knowledge, no other approach involving readily available histological specimens provides a similarly integrated picture of a disease process; moreover, it is applicable to any fixed tissue and thus, to a wide variety of normal and abnormal physiologies.

## ACKNOWLEDGEMENTS

We thank David Weinstock, Raquel Arias-Camison, Linda Wrijil, Alyce Chen, and the Gygi Lab for scientific advice and technical assistance and NanoString Inc. for GeoMx DSP data acquisition via its Technology Access Program.

## Author Contributions

MK and ZM designed this study; MK developed the proteomic method; AJN performed quantitative data analysis; IHS, RFP AJM, LW and SS performed the pathology studies and reviewed images; MK, RJE, GAB and JM collected and/or analyzed mass spectrometry data; ZM, YAC, RJP, CJ collected and analyzed multiplexed image data; DHB, SS and PKS supervised the research; MK, ZM, AJN and PKS wrote the manuscript which was read and approved by all authors.

## Data and materials availability

Proteomics raw data and search results were deposited in the PRIDE archive^56^ and can be accessed under ProteomeXchange accession numbers: PXD021453, PXD021456 and PXD021459. Datasets will be publicly available when paper is published. No unique reagents were generated by this study.

## ONLINE METHODS

### FFPE proteomics

Tissue from de-waxed formalin-fixed paraffin-embedded (FFPE) samples was scraped off slides using a fresh razor blade for each slide. Visible major blood vessels were avoided. The samples were transferred to individual PCR tubes with 100µL lysis buffer (50mM Tris pH8.5, 2% SDS, 150mM NaCl). Disulfide bonds were reduced by 5mM dithiothreitol (DTT). Crosslinks were thermally reversed by heating to 90°C for 10 minutes, sonicating for 3 minutes in a bath sonicator (Branson), and additional heating at 90°C for 10 minutes. Samples were then cooled to room temperature and diluted to 200µL with lysis buffer. Reduced cysteines were alkylated by adding freshly prepared 0.4M iodoacetamide (IAA) in 50mM ammonium bicarbonate to obtain a final concentration of 25mM IAA for 25 minutes in the dark and quenched with 1M DTT to obtain a final concentration of 50mM DTT.

Samples were transferred to clean 1.5mL microcentrifuge tubes and precipitated with a 1:1 volume ratio of 55% ice-cold trichloroacetic acid (TCA) during incubation on ice for 15 minutes. Samples were then centrifuged at ∼20,000 x g at 4°C for 10 minutes and the supernatant was aspirated. The pellets were washed with cold acetone (−20°C), which was immediately aspirated, and washed three more times with −20°C cold acetone followed by centrifugation at ∼20,000 x g at 4°C for 10 minutes. The pellet was solubilized by vortexing in 25µL of 8M urea in 200mM EPPS pH 8.5. Samples were diluted with 200 mM EPPS pH 8.5 and 4% v/v acetonitrile (ACN) buffer to achieve a final urea concentration of 1M. The protein samples were digested with Lysyl Endopeptidase (LysC) (40mg/ml, Wako, 129-02541) for 3 hours at 37°C. The sample was diluted to 0.5M urea with 200mM EPPS pH8.5 and further digested with trypsin (Promega, V5111) at a 1:100 sample/stock solution (v/v) ratio for 8 hours at 37°C. The digests were centrifuged at ∼20,000 x g for 5 minutes and supernatants were transferred to clean microcentrifuge tubes.

Acetonitrile (ACN) was added to each sample to a concentration of 30% and peptides were labeled with TMTpro reagents (Thermo Fisher Scientific, A44520) for 1 hour at room temperature with brief mixing by vortexing every 10 minutes. After labeling, a ratio check was performed by pooling 2.5µL of each reaction and label incorporation (>95%) and ratios were determined by mass spectrometry. The reactions were then quenched with a final concentration of 0.5% hydroxylamine for 15 minutes, samples mixed into a single 16-plex, ratios determined by MS analysis again and the sample acidified with formic acid (FA) to a final concentration of 2% v/v. Labeled samples were pooled to obtain on average a 1:1 ratio of TMT-labeled peptides The peptide pool was dried by SpeedVac, reconstituted in 1% FA with 0.1% trifluoroacetic acid (TFA) and desalted using a Sep-Pak C18 cartridge (200mg, Waters, WAT054925) to remove salt. The cartridge was pre-conditioned with 1% FA with 0.1% TFA and the sample was slowly loaded by gravity flow followed by 2 washes with aqueous 1% FA. Peptides were eluted with 85% ACN with 15% aqueous 1% FA.

Once dried by SpeedVac, samples were phospho-enriched using High-Select(tm) Fe-NTA Phosphopeptide Enrichment cartridges (Thermo, A32992). The flow-through was retained for further analysis. The phospho-enriched sample was desalted by C18 Empore Extraction Disks (Fisher Scientific, 13-110-019) solid-phase exchange (Stage Tip) and dried. The final phospho-enriched peptide sample was reconstituted in aqueous 1% FA without ACN for MS analysis.

The retained flow-through was purified by Sep-Pak as above again with binding to the column in 1% FA without TFA and an intermediate washing step of 1% FA with 5% acetonitrile to remove unincorporated TMTpro labels. Alkaline reversed-phase peptide fractionation was performed on an Agilent 1200 system using a flow rate of 600µL/minute and 18-45% Buffer B over 75 minutes. (Buffer A is 10mM ammonium bicarbonate pH 8 with 5% ACN, Buffer B is 10% 10mM ammonium bicarbonate pH 8 with 90% ACN). A total of 96 fractions was collected and combined into 24 samples, which were dried, desalted by stage tipping, and dried. The fractions were reconstituted in aqueous 1% FA with 3% ACN prior to MS analysis.

### Mass spectrometry analysis

All samples were analyzed using an Orbitrap Lumos or Eclipse Tribrid MS with an EASY-nLC 1200 system (Thermo Fisher Scientific) with a capillary column (length: 30cm, inner diameter: 75mm) packed with 2.6µm Accucore C18 resin and heated to 60°C. Phospho-enriched peptides were analyzed with multi-stage activation to account for phospho neutral loss (−97.977/z) and gradients ranging from 3-95% acetonitrile. Deep fractionated samples were separated with a gradient of 5-95% ACN in 0.1% FA over 5 hours and a flow rate of 250nL/minute and analyzed by MS with a MultiNotch MS^3^ method.^21^ MS^1^ scans were performed in the Orbitrap using a scan range of 400-1400m/z and dynamic exclusion of 5 minutes. MS/MS scans were performed in the Ion Trap with 35% collision energy and injection times of up to 300ms. The TMTpro reporter ions were quantified in the Orbitrap using synchronous precursor selection (SPS-MS^3^) with a scan range of 100-1000m/z and an HCD collision energy of 40%. The Orbitrap resolution was set to 50,000 (dimensionless units) and the maximum injection time varied with up to 650ms. Further details on TMTpro multiplexing, LC and MS parameters and settings were identical to those recently described.^57^

An in-house developed software suite was used to convert recorded.raw files to mzXML format, correct monoisotopic m/z measurements, and perform a post-search calibration. Peptide-spectrum matches used a Sequest (v.28, rev.12) based search engine. Searches were performed using a mass tolerance of 20ppm for precursors and a fragment ion tolerance of 0.9Da against databases with the *Macaca mulatta* reference proteome (UniProt) with additional *Macaca mulatta* proteins or a human reference proteome (2018, UniProt) and SARS-CoV-2 proteins (UniProt). Search databases contained common contaminant proteins and reversed sequences as decoys. Search results were filtered by linear discriminant analysis (LDA) against reversed peptide sequences with a false discovery rate (FDR) set to <1% and a protein FDR <1% for collapsed protein groups. Search parameters accounted for oxidized methionine (+15.9949Da) dynamically with TMTpro label (+304.2071Da) on peptide N-termini and lysine as static modification and in the case of phospho-peptide samples dynamic phospho-Ser/Thr/Tyr (+79.9663Da). A modified Ascore algorithm was applied to assess the confidence of phosphorylation-site position assignment.^58^ Phosphorylation localized to individual sites required Ascore values >13 (p ≤0.05) for confident localization. Site localization score (Max Score) for phosphopeptides quantified is reported in Supplementary Table 5. MS^3^ was used to quantify relative TMTpro signal to noise (s/n), peptide quant data were filtered for an isolation specificity of greater than 70% and a sum s/n of greater than 200 over all sixteen channels for each peptide. Relative protein quantification was achieved by calculating the summed s/n for all peptides mapping to a given protein. Channels were normalized for equal total protein content by summed s/n for all proteins in a given channel. Details of the TMT intensity quantification method and further search parameters applied were described previously.^59^ Proteomics raw data and search results were deposited in the PRIDE archive^56^ and can be accessed under ProteomeXchange accession numbers: PXD021453, PXD021456 and PXD021459 (note to reviewers: datasets will be publicly available when paper is published, per PRIDE policy. Data are available during the review phases at https://www.ebi.ac.uk/pride/login with Username, Password: LPCv6VLi; Username, Password: mJoqEZJU and Username, Password: 1ruKkhd5).

### Cross-linking analysis

To identify transglutaminase cross-linked fibrin peptides, all MS data were searched with the PIXL search engine,^60^ specifying the Lys - Gln residue linkage alongside regular peptide search. Fibrinogen and fibronectin proteins were included in cross-linked search, along with serum albumin, alpha-2-macroglobulin and alpha-2-antiplasmin – all proteins previously implicated with transglutaminase activity.^61^ Precursor tolerance was set to 15ppm and fragment ion tolerance to 0.4Th. Due to the low number of cross-linked peptides in the sample, they were filtered to 1% FDR using LDA alongside regular peptides.

### Statistical analysis of MS data

Relative summed s/n TMT data for 16-plex experiments were scaled (0-100%) for each protein and analyzed by principal component analysis (PCA) and 2-way hierarchical clustering (Ward) using the JMP Pro 15 software package. DESeq2^62^ was also used for differential expression analysis. A corrected P-value cut-off of 0.05 was used to assess significant genes that were up-regulated or down-regulated after application of the Benjamini-Hochberg procedure.^63^

### Pathway enrichment analyses

A compendium of biological and immunological signatures was identified from publicly available databases or published manuscripts for performing enrichment analysis. In order to perform gene set enrichment analysis, two previously published methods (Gene Set Enrichment Analysis (GSEA)^64^ and Single-sample GSEA (ssGSEA) were primarily used. The R package clusterProfiler^65^ was used to perform GSEA analysis and the R package GSVA^66^ was used to perform ssGSEA analysis which calculates the degree to which genes in a particular gene set are coordinately up- or down-regulated within a sample. The interferon, complement, platelet, fibrin, IL1, IL1R, NF-KB, IL10, MTORC and insulin related signatures were curated from MSigDB^67^ and immune cell related signatures were manually curated from published studies^68–70^.

### Cyclic immunofluorescence

Tissue-based Cyclic Immunofluorescence (t-CyCIF) was performed as previously described^20^ with antibody validation as described^71^. The method involves iterative cycles of antibody incubation, imaging, and fluorophore inactivation. Briefly, FFPE specimens on glass slides were de-waxed and antigen retrieval was performed on a Leica Bond RX. Each section of tissue was stained and imaged with fluorescently conjugated antibodies or unconjugated primary and conjugated secondary antibody (Table S6). Slides were scanned with a RareCyte CyteFinder scanner equipped with a 20x, 0.75NA objective. The raw images were corrected for uneven illumination using the BaSiC tool^72^ and images from multiple rounds of scanning of the same tissue were stitched and aligned using ASHLAR.^73^. Cell-nucleus pixel probability maps were first generated using a trained U-Net^74^ model and then single cells were segmented using marker-controlled watershed. Mean fluorescence intensities of each marker for each cell were computed to determine cell types. In Fig. 2-4 selected CyCIF channels are shown

### GeoMx analysis

NanoString GeoMx was performed as previously described^75^ at NanoString Inc. as part of their DSP Technology Access Program. 5μm sections from high-virus day 2 and uninfected NHP tissue blocks were used. Immunofluorescence with a DNA stain and antibodies against PANCK, CD45 were used to visualize tissue, with area of interest (AOI) guided by staining of SARS-N and DNA staining in an adjacent stained tissue section. 67 AOIs representing five morphological sites (uninfected/control (24), airway (2), two virus positive regions (25) with differential levels of immune infiltration as well as a virus-negative region of consolidated region (16)) were extracted and assayed. The sequencing and probe quality-controlled data received directly from NanoString were then used for pathway analysis.

## EXTENDED DATA FIGURES

**Extended Data Figure 1.**
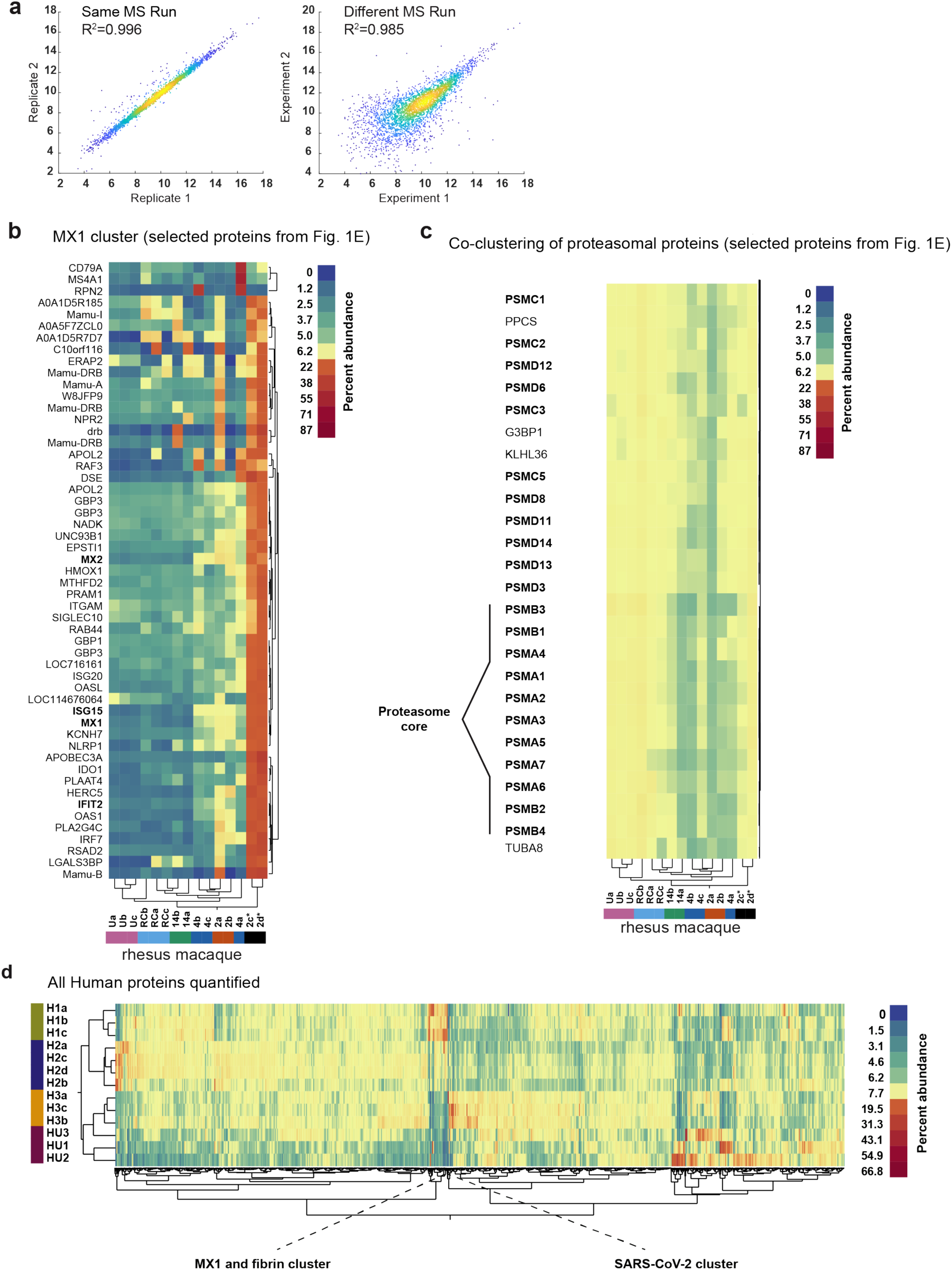
Reproducibility of quantitative proteomics data and trajectory of antiviral response to SARS-CoV-2 infection. **a**, left: Data reproducibility between two technical replicates (two FFPE samples from the same specimen) in which TMT labeling and MS analysis were performed in parallel. Right: two adjacent FFPE sections from the same tissue block processed on different days and then labelled with TMT and subjected to MS separately. Comparing similar samples between different multiplexes, peptides (proteins) having low TMT summed signal-to-noise ratios scatter around the trend line. **b**, MX1 cluster, magnification of Fig. 1f. Antiviral immune response factors (MX1, ISG15, MX2 and IFIT2 as examples in bold) closely cluster together in two-way hierarchical cluster analysis. Please note that in virus positive specimens (2c-d, black) all proteins of the MX1 cluster display a strong increase over other specimens from infected animals and uninfected control specimens. Antiviral response proteins decrease from virus positive 2 days post infection samples (d.p.i., black) over virus low day 2 (orange) and 4 (dark blue) samples to 14 (dark green) samples. In viral re-challenge specimens (re, blue) antiviral proteins levels are largely at uninfected (U, pink) baseline. Scaled relative protein abundance range from 0% (dark blue) to 87% (dark red). **c**, Proteasome cluster magnification of Fig. 1e. Proteasomal proteins (bold) cluster together in quantitative mass spectrometry data. Alpha (PSMA1-7) and beta subunits (PSMB1-4) of the proteasome core co-cluster within this group. Scale as in c. **d**, Two-way hierarchical clustering of 6,729 quantified proteins from 13 human FFPE lung specimens of 3 SARS-CoV-2 infected (H1, green, H2, navy blue, H3, yellow, respectively and in 3 uninfected (HU, red) decedents. Samples group by infection status. Positions of the MX1 and fibrin protein cluster and the viral S and N protein cluster are indicated. MX1/fibrin and SARS-CoV-2 clusters are separated by the first dendrogram branch of the full dataset. Scaled relative protein abundance range from 0% (dark blue) to 66.8% (dark red).

**Extended Data Figure 2.**
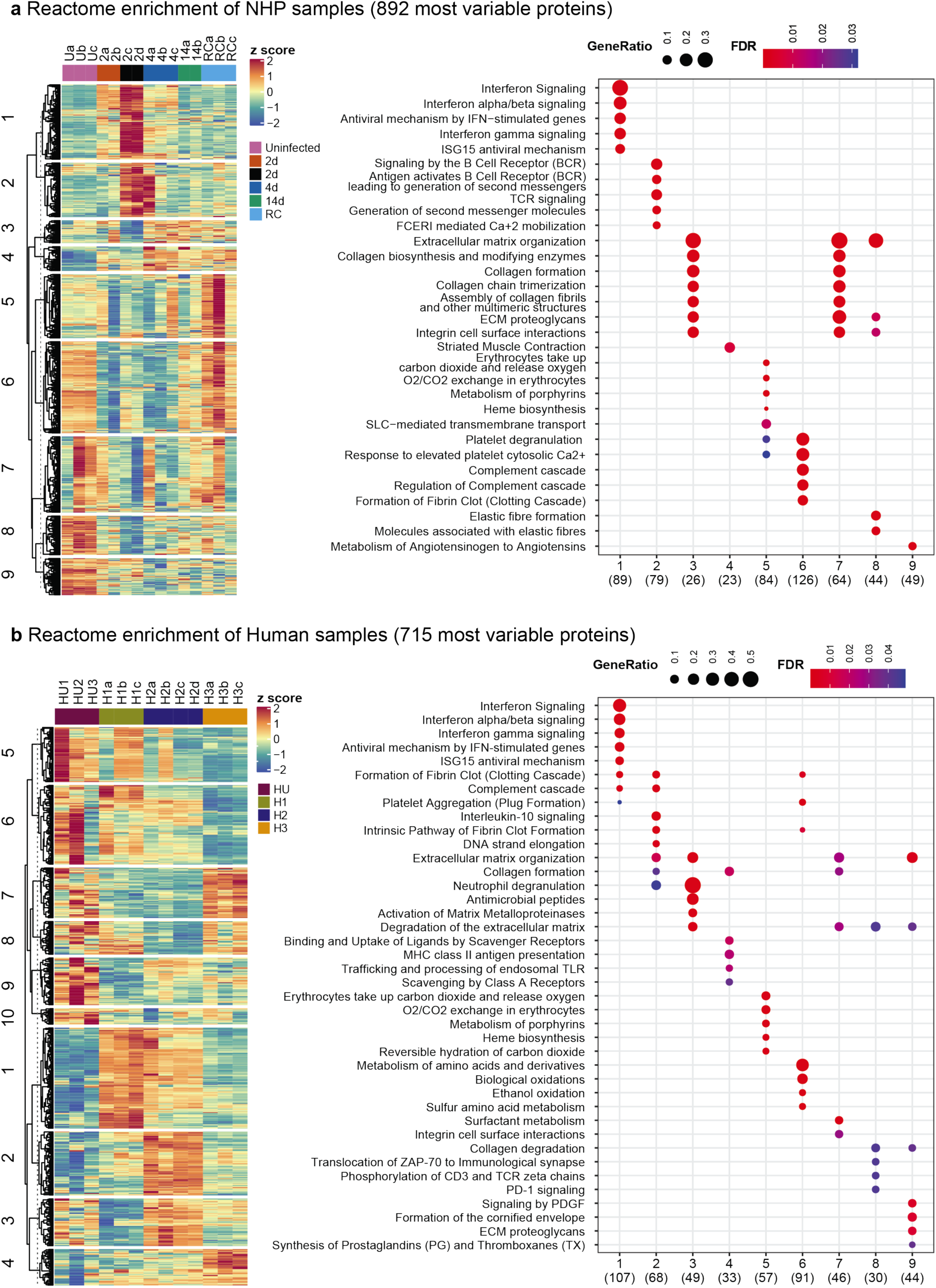
Reactome enrichment analysis of the most variable proteins in NHP and Human samples. **a**, Heatmap of 892 most variable proteins divided into 9 clusters by k-Means clustering (left panel) and the corresponding top 10 Reactome enrichment terms (right panel; FDR <0.05). **b**, Heatmap of 715 most variable proteins divided into 10 clusters by k-Means clustering (left panel) and their corresponding top 10 Reactome enrichment terms (right panel; FDR <0.05). Protein gene ratios range from 0.1 to 0.3 in NHPs and 0.1 to 0.5 in humans (scaled marker size).

**Extended Data Figure 3.**
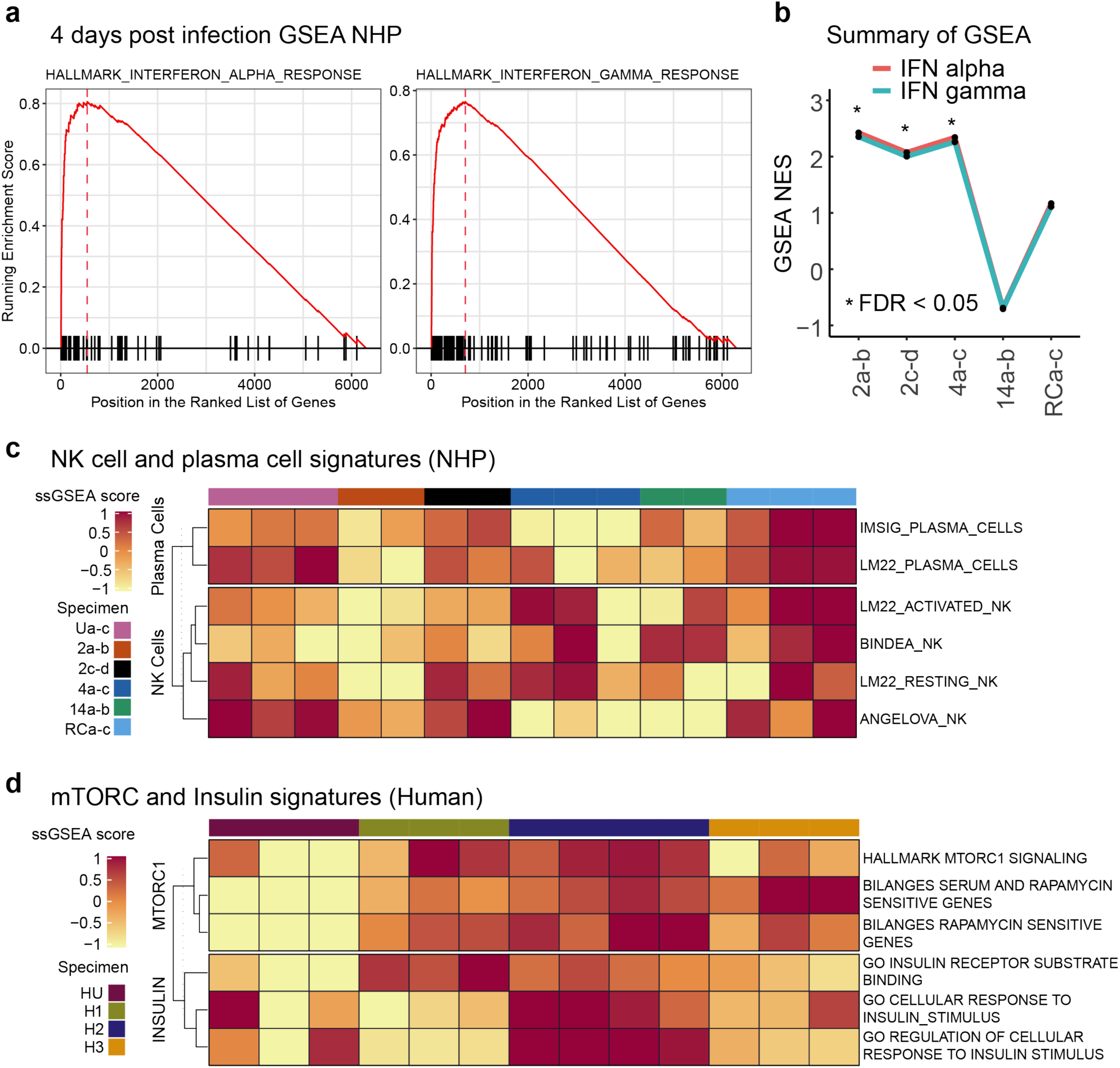
Analysis of Interferon-stimulated genes (ISGs) and antiviral proteins of SARS-CoV-2 challenged non-human primate (NHP) specimens and clinical human decedent samples. **a**, Gene set enrichment analysis (GSEA) plot for hallmark IFNα (left panel) and IFNγ (right panel) (FDR <0.05). The red plot corresponds to the running enrichment score for the pathway as the analysis walks down the ranked list, including the location of the maximum enrichment score (peak of the curve). The small black bars on the x-axis show where the members of the gene set appear in the ranked list of genes. The plot presents data from NHP samples at day 4 post infection compared to uninfected samples. **b**, Line plot summarizes enrichment scores from GSEA of hallmark IFNα and IFNγ signatures for each time point post infection in NHP samples compared to uninfected samples (* FDR <0.05). **c-d**, ssGSEA heatmap showing the enrichment of NK and plasma cell and mTOR and Insulin response **d**, related gene signatures in NHP samples. The enrichment scores are normalized by the absolute difference between minimum and maximum values and scaled between −1 to 1.

**Extended Data Figure 4.**
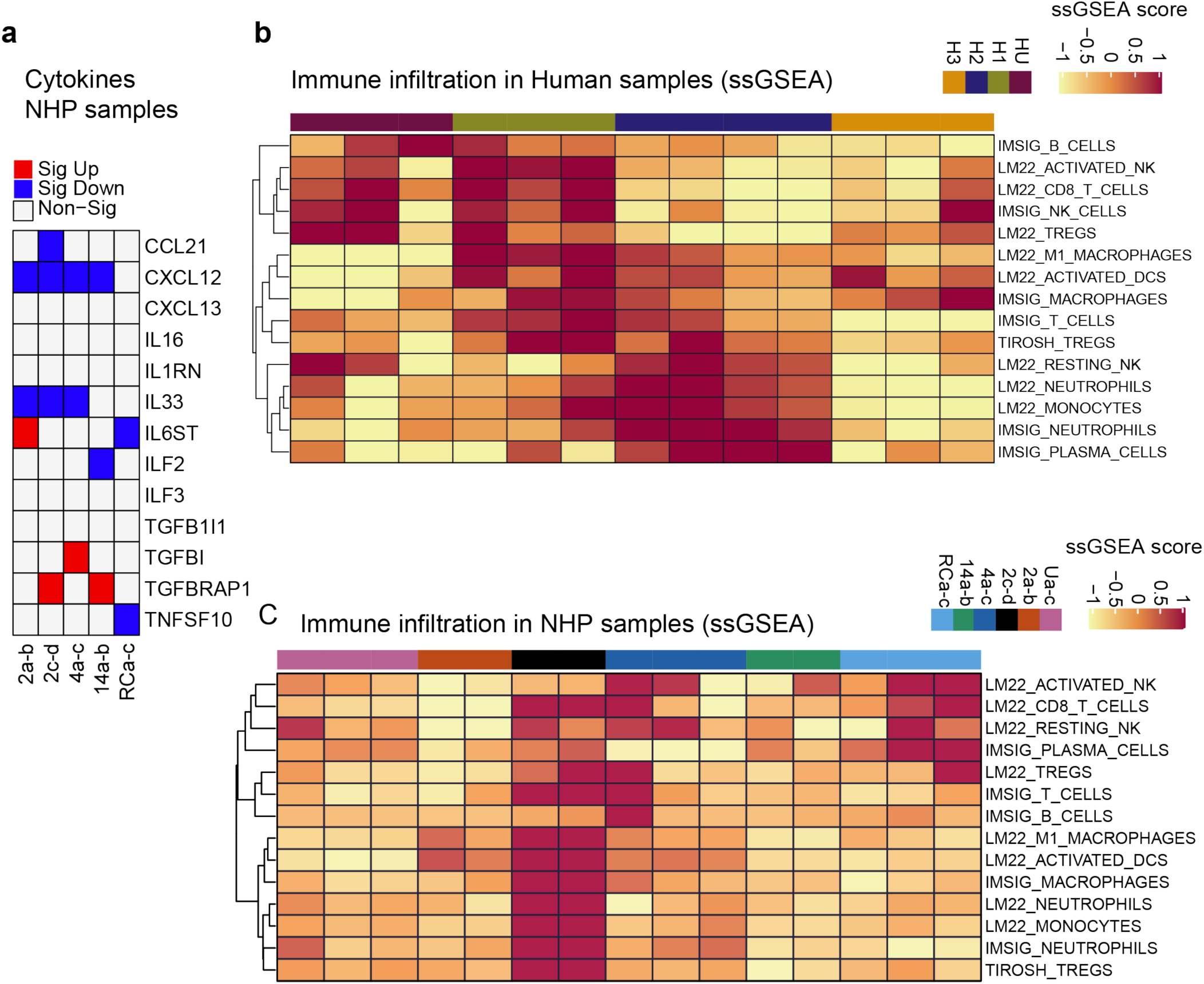
Enrichment analysis of cytokine levels and inflammation in FFPE lung samples from NHP and human decedents. **a**, Binary heat map showing significant increase (red) or decrease (blue) of quantified cytokines in infected relative to uninfected NHP samples by mass spectrometry (P < 0.05, t-test). **b**, ssGSEA heatmap showing the enrichment of immune cell related gene signatures in human decedents. The enrichment scores are normalized by the absolute difference between minimum and maximum and scaled between −1 to 1. **c**, ssGSEA heatmap as in panel B but for NHP samples.

