## Supplementary Table 1 for "Multiplexed proteomics and imaging of resolving and lethal SARS-CoV-2 infection in the lung"

**Table S1: Characteristics and identities of non-human primate specimens**

| sample | group | pathogen | location | animal | block | time | MS | Imaging |
|---|---|---|---|---|---|---|---|---|
| Ua | Control | uninfected | unknown | 5841 | none | Control | Yes | Yes |
| Ub | Control | uninfected | unknown | 5841 | none | Control | Yes | Yes |
| Uc | Control | uninfected | unknown | 5841 | none | Control | Yes | Yes |
| Ud | Control | uninfected, fetal | unknown | 583 | none | Control | No | Yes |
| 2a | 2 day virus neg | SARS-CoV2 | Right Cranial 4_5 | 7255 | DB01_6 | 2 day | Yes | Yes |
| 2b | 2 day virus neg | SARS-CoV2 | Right middle_6 | 7253 | DB01_7 | 2 day | Yes | Yes |
| 2c* | 2 day virus pos | SARS-CoV2 | Right Caudal_8 | 7255 | DB01_8 | 2 day | Yes | Yes |
| 2d* | 2 day virus pos | SARS-CoV2 | Right (most likely cranial) | 7255 | DB01_30 | 2 day | Yes | Yes |
| 2e | 2 day virus pos | SARS-CoV2 | Right (most likely cranial) | 7253 | DB01_9 | 2 day | No | Yes |
| 4a | 4 day | SARS-CoV2 | unknown | 7260 | DB01_5 | 4 day | Yes | No |
| 4b | 4 day | SARS-CoV2 | Right Cranial_5 | 7260 | DB01_6 | 4 day | Yes | Yes |
| 4c | 4 day | SARS-CoV2 | Left cranial_3 | 7259 | DB01_9 | 4 day | Yes | Yes |
| 4d | 4 day | SARS-CoV2 | Left cranial_3 | 7259 | DB01_7 | 4 day | No | Yes |
| 14a | 14 day | SARS-CoV2 | Right Cranial_3 | BS80 | DB11_1 5 | 14 day | Yes | Yes |
| 14b | 14 day | SARS-CoV2 | Right Cranial_3 | 7154 | DB11_2 5 | 14 day | Yes | Yes |
| RCa | 35 day rechallenge | SARS-CoV2 | Right Cranial_3 | 7162 | DB05_5 | 35 day rechallenge | Yes | Yes |
| RCb | 35 day rechallenge | SARS-CoV2 | Right Cranial_3 | 7173 | DB05_5 | 35 day rechallenge | Yes | Yes |
| RCc | 35 day rechallenge | SARS-CoV2 | Right Cranial_3 | 7244 | DB05_5 | 35 day rechallenge | Yes | Yes |
