## Supplementary Table 3 for "Multiplexed proteomics and imaging of resolving and lethal SARS-CoV-2 infection in the lung"

### Table S3: Characteristics of human COVID-19 decedents and controls

| Specimen Code | Age | Sex | Notable medical history | Time to death from admission | Post-mortem interval | Cause of death | Lung weights at autopsy* | Lung histology at autopsy | Other autopsy findings |
|---|---|---|---|---|---|---|---|---|---|
| **H1a-c** | 66 | F | SLE<br>Rheumatoid arthritis<br>Pulmonary fibrosis<br>Chronic kidney disease<br>Interstitial lung disease<br>MGUS<br>Coronary artery disease<br>Systemic hypertension | 7 | 2 | Respiratory failure | Right 900g; Left 630g | Diffuse alveolar damage<br>Interstitial lung disease with bronchiectasis | Cardiomegaly (620g) |
| **H2 a-d** | 57 | M | Systemic hypertension<br>Type 2 diabetes mellitus | 1 | 1 | Respiratory failure | Right 1230g; Left 1210g | Diffuse alveolar damage | Cardiomegaly (460g) |
| **H3a-c** | 68 | F | CAD<br>Atherosclerosis<br>Systemic hypertension<br>Type 2 diabetes mellitus<br>COPD | 1 | 1 | Cardiac/respiratory failure | Right 900g; Left 380g | Diffuse alveolar damage | Acute myocardial infarction<br>Ischemic bowel<br>Cardiomegaly (680g) |
| **HU1** | 51 | M | CAD<br>Hypertension<br>Hyperlipidemia<br>Obesity<br>GERD<br>OSA | <1 | 1 | Atherosclerotic CAD | Right 720g; Left 460g | No significant pathologic change | Myocardial infarction<br>Acute myocardial ischemia<br>Cardiomegaly (600g) |
| **HU2** | 75 | F | Poorly-differentiated infiltrating ductal carcinoma | 0 | 2 | Sudden cardiac death suggesting arrhythmia | Right 500g; Left 500g | Pulmonary parenchyma with mild-moderate emphysema | Systemic atherosclerosis<br>Valvular heart disease |
| **HU3** | 51 | M | Liver cirrhosis | | 4 | Hepatorenal failure | - | No significant pathologic change | Acute steatohepatitis<br>Acute renal failure<br>Acute pneumonia<br>Colitis |

*Reference range: Female: Right 360-570g, Left 325-480g; Male: Right 360-570g, Left 325-480g
