## Supplementary Table 6 for "Multiplexed proteomics and imaging of resolving and lethal SARS-CoV-2 infection in the lung"

**Table S6**

| Target | Fluorochrome | Species | Clone | Vendor | Catalog No | dilution_H | dilution_NHP | Concentration (µg/ml) | reactivity |
|---|---|---|---|---|---|---|---|---|---|
| aSMA | eFluor 660 | Mouse | 1A4 | Thermo Fisher | 50-9760-82 | 1000 | n.a. | n.d. | H, NHP |
| aSMA | Alexa Fluor 750 | Mouse | 1A4 | R&D | IC1420S | n.a. | 300 | n.d. | H, NHP |
| CD16 | Alexa Fluor 647 | Mouse | DJ130c | Santa Cruz Biotechnologies | sc-20052 AF647 | 150 | 150 | n.d. | H, NHP |
| CD163 | Alexa Fluor 488 | Rabbit | EPR14643 | Abcam | ab218293 | 300 | n.a. | n.d. | H |
| CD163 | none | Mouse | 10D6 | Thermo Fisher | MA5-11458 | n.a. | 50 | n.d. | H, NHP |
| CD206 | Alexa Fluor 488 | Mouse | D-1 | Santa Cruz Biotechnologies | sc-376108 AF488 | n.a. | 100 | n.d. | H, NHP |
| CD31 | Alexa Fluor 647 | Rabbit | EPR3094 | Abcam | ab218582 | 200 | 200 | n.d. | H, NHP |
| CD61 | Alexa Fluor 647 | Mouse | VI-PL2 | BioLegend | 336408 | 200 | n.a. | n.d. | H |
| CD68 | Phycoerythrin | Rabbit | D4B9C | CST | 79594S | 200 | 300 | n.d. | H, NHP |
| HLADR | Alexa Fluor 555 | Rabbit | EPR3692 | abcam | ab215312 | 400 | 400 | n.d. | H, NHP |
| IFIT3 | Alexa Fluor 647 | Mouse | B-7 | Santa Cruz Biotechnologies | sc-393512 AF647 | n.a. | 150 | n.d. | H, NHP |
| KRT7 | Alexa Fluor 555 | Rabbit | EPR17078 | abcam | ab209601 | 1000 | 2000 | n.d. | H, NHP |
| Mouse IgG | Alexa Fluor 647 | Goat | polyclonal F(ab') | Thermo Fisher | A21237 | 2000 | 2000 | 0.1 | H, NHP |
| MPO | Alexa Fluor 488 | Rabbit | EPR20257 | Abcam | ab225474 | 400 | 300 | n.d. | H, NHP |
| MX1 | Alexa Fluor 488 | Rabbit | D3W7I | CST | 79373BC | n.a. | 150 | n.d. | H, NHP |
| Rabbit IgG | Alexa Fluor 488 | Goat | polyclonal F(ab') | Thermo Fisher | A11070 | 2000 | 2000 | 0.1 | H, NHP |
| SARS-N | none | Rabbit | polyclonal | Novus | NB100-56576 | 250 | 300 | n.d. | H, NHP |
| VIM | Alexa Fluor 555 | Rabbit | D21H3 | CST | 9855S | n.a. | 300 | n.d. | H, NHP |
